# Establishing a transcriptome-based drug discovery paradigm for neurodevelopmental disorders

**DOI:** 10.1101/2020.05.13.093468

**Authors:** Ryan S. Dhindsa, Anthony W. Zoghbi, Daniel K. Krizay, Chirag Vasavda, David B. Goldstein

## Abstract

Advances in genetic discoveries have created substantial opportunities for precision medicine in neurodevelopmental disorders. Many of the genes implicated in these diseases encode proteins that regulate gene expression, such as chromatin associated proteins, transcription factors, and RNA-binding proteins. The identification of targeted therapeutics for individuals carrying mutations in these genes remains a challenge, as the encoded proteins can theoretically regulate thousands of downstream targets in a considerable number of cell types. Here, we propose the application of a drug discovery approach called “transcriptome reversal” for these disorders. This approach, originally developed for cancer, attempts to identify compounds that reverse gene-expression signatures associated with disease states.

## Introduction

The introduction of whole-exome sequencing has led to genetic discoveries that have dramatically improved our understanding of mechanisms underlying neurodevelopmental disorders^1^. Some of these discoveries have translated to the development of targeted treatments^2^, including enzyme replacement therapies in deficiency disorders^3^, antisense oligonucleotide therapies for spinal muscle atrophy^4,5^, and channel modulators for gain-of-function channelopathies^6^. Despite these important achievements, the development of precision therapeutics has lagged far behind the rapid progress in gene discovery.

One of the major challenges facing precision medicine in neurodevelopmental disorders—including autism spectrum disorder (ASD), developmental epileptic encephalopathy (EE), developmental delay with cognitive manifestations (DD), and schizophrenia—is the tremendous genetic heterogeneity underlying these conditions. Implicated genes have revealed a wide range of etiologies for each of these diseases, such as abnormal synaptic transmission, chromatin remodeling, transcription regulation, and ion channel function^7–11^. It is likely that this heterogeneity will require different drug discovery paradigms for each functional class of genes.

We argue that neurodevelopmental disorder associated genes that directly influence the transcriptome—namely chromatin modifiers, transcription factors and co-factors, and RNA-binding proteins—represent one particular group of genes amenable to drug discovery efforts that are well-developed in cancer but largely unexplored in neurology. This strategy, termed transcriptome reversal, posits that if gene expression changes underlie the pathophysiology of a particular disease, then correcting this transcriptomic signature toward a normal state may have therapeutic potential^12,13^.

Here, we outline a strategy for the development of a systematic program for transcriptome-guided drug discovery in neurodevelopmental disease. We first provide an overview of the transcriptomic signature reversal approach and its successes in other disease areas. We next discuss opportunities for analogous approaches in neurodevelopmental disorders by focusing on transcriptional regulators that have been implicated in EE, ASD, DD, and schizophrenia. Finally, we propose immediate next steps that would be required to systematically implement this drug discovery strategy.

### The transcriptome reversal paradigm

In theory, many different classes of molecules can reverse transcriptomic signatures toward a healthy state, including small-interfering RNAs, antisense oligonucleotides, peptides, and classical small molecules. Here we focus on small molecules because they are especially pharmacologically versatile and have been the focus of most transcriptome reversal efforts to date. Broadly speaking, this approach requires three steps: identification of the disease gene expression signature, *in silico* or experimental screening to prioritize compounds most likely to reverse this disease signature, and targeted experimental validation of candidate compounds (**Figure 1**).

**Figure 1.**
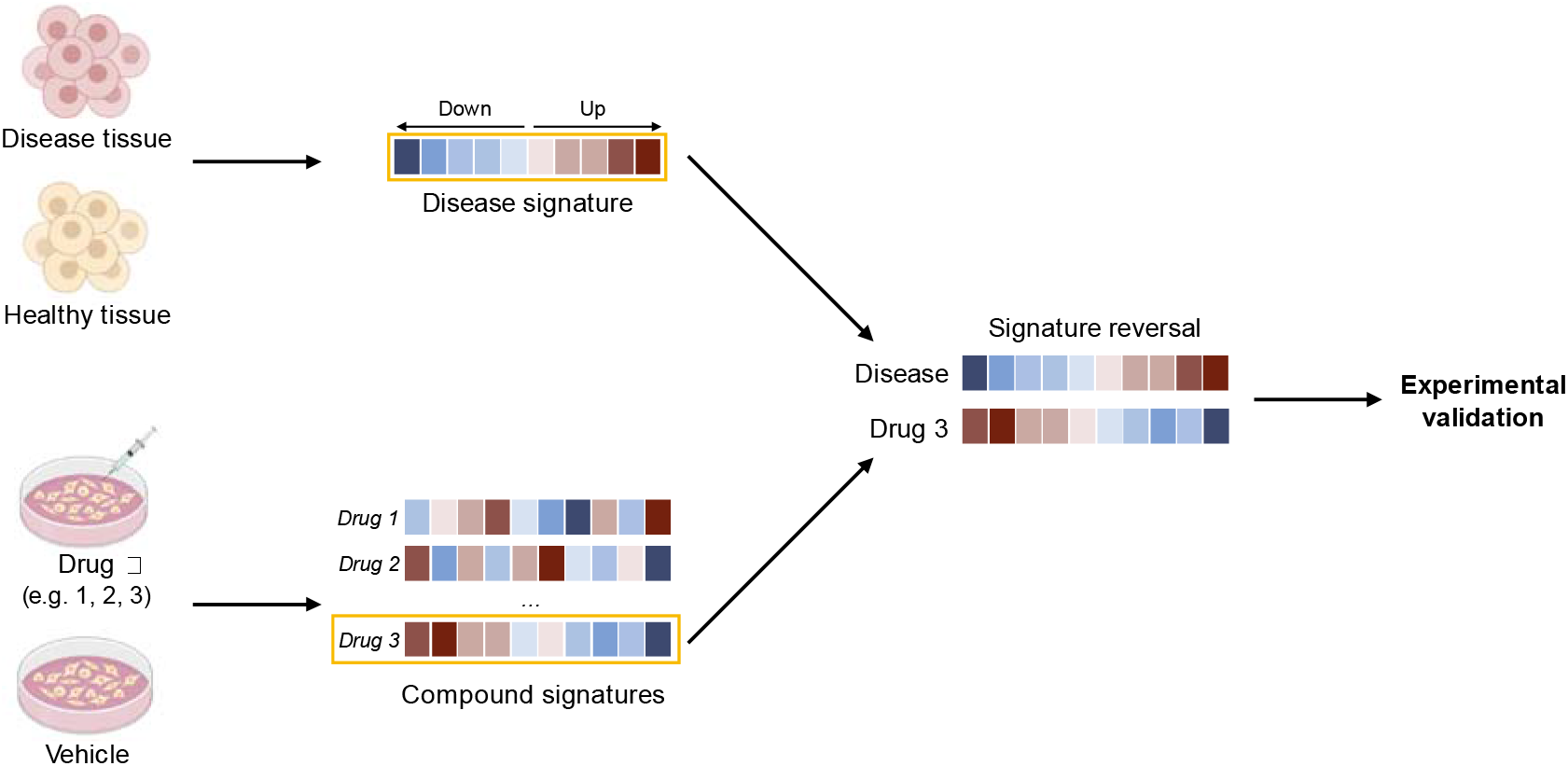
The transcriptome reversal approach. Gene expression profiling (e.g. RNA-seq or single-cell RNA-seq) is used to derive disease expression signatures. This signature is then compared to the signatures of cells treated with small molecules in order to identify compounds most likely to reverse disease-associated expression changes. Candidate compounds are then screened experimentally to determine whether they reverse disease phenotypes.

### Generating disease gene expression signatures

Generating disease expression signatures requires the unbiased assessment of gene expression changes in tissue or cells derived from patients or a disease model. By performing RNA-sequencing on disease samples and healthy samples, researchers can then use differential gene expression analysis (DGE) to identify quantitative gene expression level changes between sample groups. This strategy reveals the sets of genes that are up- and down-regulated in the disease state compared to a normal state (i.e. the disease expression signature). In addition to being useful for transcriptome reversal, these expression signatures can reveal perturbed biological pathways and provide insight into disease mechanisms^14^.

Importantly, advances in single-cell RNA-sequencing (scRNA-seq) allow interrogation of these expression changes at an unprecedented resolution. Until recently, RNA-seq was predominantly performed on bulk tissues. While these studies led to transformative discoveries, bulk RNA-sequencing methods cannot resolve specific cell types and only provide an average expression signal for an ensemble of cells. scRNA-seq, on the other hand, can resolve cell type-specific expression signatures. The most common scRNA-seq methods involve microfluidic chips that encapsulate single cells into reagent-filled oil droplets^15^. Reverse transcription occurs within each oil droplet, and the resulting cDNAs receive barcodes that allow for the assignment of each resulting sequencing read back to its own single cell of origin. It is important to note that performing scRNA-seq necessitates particularly careful experimental design and analyzing the resulting data requires statistical rigor^14,16^.

### *In silico* screening of small molecules

Once a disease gene expression signature has been identified, the next step is to identify compounds that are predicted to reverse these expression changes. This process requires two components: (1) a database of gene expression signatures of cells or tissues treated with small molecules; and (2) a statistical method to compare the disease signature to these small molecule signatures.

There have been considerable efforts to create publicly available compendia of small molecule expression signatures. In 2006, Lamb and colleagues introduced the “Connectivity Map” (CMap), which included microarray-derived signatures for roughly 1,300 small molecules applied to human cell lines^13^. More recently, a new version of the Connectivity Map was introduced that leverages a novel gene expression profiling technique called L1000^17^. This assay measures the expression level of 978 “landmark genes” that were selected to capture a large proportion of genome-wide variation in gene expression. The measurements of these genes are then used to infer the expression of 11,350 other genes in the transcriptome. Because the L1000 assay is far more affordable than typical RNA-sequencing, CMap investigators were able to use this platform to profile roughly 20,000 small molecules in a variable number of human cell lines. This resource has not only facilitated drug discovery in cancer^18–20^, but also in other non-neurological diseases too, including diabetes^21^, inflammatory bowel disease^22^, and osteoporosis^23^ (reviewed by Keenan et al.^24^).

Investigators have introduced several statistical methods that compare disease and compound signatures to predict which compounds are most likely to invert the disease signature. For example, the CMap uses the Connectivity Score: a normalized similarity metric based on the weighted Kolmogorov-Smirnov enrichment statistic. Other methods rely on the organization of expression data into networks to infer disease- and drug-induced changes in master regulator activity^18,25^. Computational algorithms, such as OncoTreat^19^, can be used to identify individual or combinations of compounds that are expected to reverse master regulator activity in the disease state. Importantly, as discussed below, the success of this *in silico* screening step relies on the availability of cell type-specific perturbation data.

### Experimental validation

Once small molecules are prioritized via *in silico* screening, the next step is to verify that the top-scoring compounds in fact reverse the disease profile. In cases where the small molecule signatures were generated in cell types not related to the disease of interest, it is critical to appreciate that the compounds identified may not be the best compounds in cell types relevant to the disease. Therefore, validation requires the administration of candidate compounds to patient cells or cells derived from the disease model followed by gene expression profiling. The most promising compounds should reverse disease-related expression changes, such that the signature more closely resembles the wildtype signature. Furthermore, candidate compounds should be administered at multiple doses and timepoints to pinpoint the biological conditions that lead to the strongest restoration of the transcriptome. When possible, these compounds should then be evaluated for their ability to rescue disease phenotypes in a relevant disease model.

### Transcriptome reversal opportunities in neurodevelopmental disorders

Despite the promise of this approach, there are relatively few examples of transcriptome reversal studies in neurodevelopmental disorders. Most attempts have focused on non-genetic disease models or tissue derived from patients who lack a clear Mendelian diagnosis. For example, Readhead and colleagues recently used L1000 to identify compounds that reverse transcriptomic signatures of post-mortem tissue derived from 12 individuals with schizophrenia^26^. Other studies have identified compounds that reverse disease signatures identified in mouse models of acquired epilepsy^27,28^ and in surgical tissue derived from patients with unilateral mesial temporal lobe epilepsy^29^.

These studies have provided preliminary proofs of concept, but when the underlying cause of disease is unknown, it can be challenging to decipher whether the observed disease signatures reflect causal versus compensatory gene expression changes. Thus, we argue that an important path forward is to focus on genetic models in which gene expression dysregulation represents the primary mechanism of pathogenicity. Here, we define primary transcriptomic regulators as transcription factors/co-factors, chromatin-associated proteins, and RNA-binding proteins. Strikingly, these genes encompass a substantial proportion of the genes implicated in developmental delay with cognitive impairment (13%), autism spectrum disorder (45%), developmental and epileptic encephalopathy (18%), and schizophrenia (20%) (**Figure 2**). We briefly review the functions of these gene expression regulators below and provide select examples of disease-related genes that fall into each of these classes.

**Figure 2.**
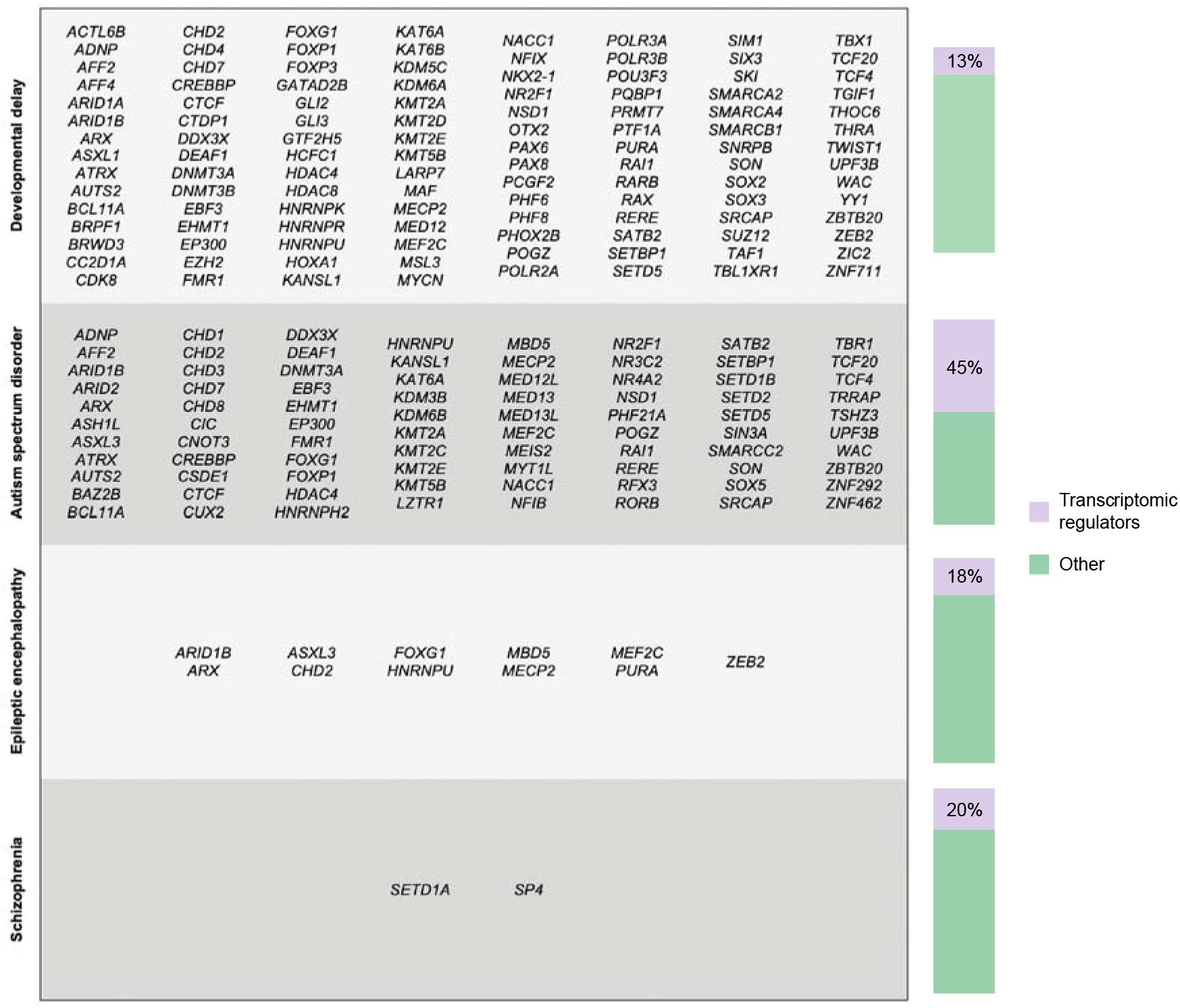
Transcriptomic regulators implicated in neurodevelopmental disorders. Table of genes encoding transcription factors, chromatin-associated proteins, and RNA-binding proteins in developmental delay with intellectual disability, autism spectrum disorder, epileptic encephalopathy, and schizophrenia. Bar charts depict the percentage of transcriptomic regulators amongst the total number of genes implicated in each disease area.

### Transcription factors

Transcription factors (TFs) bind to specific DNA motifs to recruit and regulate the transcription machinery. These proteins can be broadly separated into two classes based on their regulatory function: control of transcription initiation and control of transcription elongation^30^. This distinction is not absolute, however, as some TFs mediate both of these processes.

Furthermore, TFs can work alone or with other proteins (e.g. co-factors) in a complex to activate or repress the recruitment of RNA polymerase. Unsurprisingly, pathogenic mutations in TFs associated with neurodevelopmental disorders cause widespread transcriptomic defects in the brain.

Mutations in the transcription factor-encoding gene *MEF2C* are associated with autosomal dominant intellectual disability, autism spectrum disorder, and epilepsy^31,32^. Specifically deleting *MEF2C* in forebrain excitatory neurons in mice leads to broad changes in genes that influence neuronal specification and synapse development, leading to a dramatic increase in inhibitory synaptic transmission^33^. The FOXO (forkhead box) family of transcription factors also influences a wide network of genes important in neurodevelopment, and patients with mutations in *FOXG1* present with autism spectrum disorder^34^. Sequencing chromatin immunoprecipitation of Foxg1 identified thousands of loci in the mouse genome where this protein binds, suggesting that Foxg1 regulates hundreds of targets *in vivo*^35^.

### Chromatin-associated proteins

In eukaryotes, DNA resides in the nucleus where it is packaged into a highly condensed structure called chromatin. The primary unit of chromatin is the nucleosome, which consists of DNA wrapped around eight histone proteins. Chromatin remodelers, which move histone proteins to make DNA accessible to transcription machinery, have been implicated in neurodevelopmental disorders. Additionally, each of these histone proteins has a peptide tail that is post-translationally modified (e.g. acetylated or methylated) to activate or repress transcription. Mutations in chromatin regulators that create, remove, and recognize these chemical modifications can also result in neurodevelopmental disorders. DNA nucleotides can also be chemically modified themselves in ways that influence gene expression. The most well-studied chemical modification is DNA methylation, in which cytosines in the context of CpG dinucleotides receive methyl groups to regulate gene expression.

Loss-of-function mutations in the gene *MECP2* (methyl CpG binding protein 2), which encodes a methyl CpG binding protein, cause Rett Syndrome^36^. Conditionally ablating *MECP2* from the mouse brain recapitulates several findings in patients with Rett Syndrome, such as uncoordinated gait and repetitive movements^37,38^. Gene expression profiling of mouse models as well as human brain tissue has revealed widespread changes in gene expression across brain regions^39–41^, consistent with *MECP2’s* role in modulating chromatin topology.

### RNA-binding proteins

RNA-binding proteins play a key role in the post-transcriptional control of RNAs^42^. They regulate nearly all aspects of RNA biogenesis, including RNA splicing, polyadenylation, mRNA localization and degradation, and translation. There are hundreds of RNA-binding proteins with diverse functions, each with different RNA-sequence specificities and affinities. Many of these proteins seem to bind to thousands of transcripts per cell. Indeed, pathogenic mutations in these genes can lead to the dysregulation of hundreds or thousands of genes. For example, mutations in *HNRNPU*, an RNA- and DNA-binding protein, is associated with epileptic encephalopathy and autism spectrum disorder^7,43,44^. We recently reported that a mouse model of *HNRNPU* haploinsufficiency exhibits neuroanatomical abnormalities, global developmental delay, and increased seizure susceptibility^45^. Single-cell RNA-sequencing of mutant hippocampal and neocortical cells revealed thousands of dysregulated genes across neuronal subtypes that converged on biological pathways important to neurodevelopment^45^.

### Other gene expression regulators

Although we focus here on transcription factors/co-factors, chromatin-associated proteins, and RNA-binding proteins, we note that other genes implicated in neurodevelopmental disorders may also affect gene expression. Some examples include kinases, G-protein subunits, and ubiquitin ligases. Many of these proteins, however, are also directly involved in many alternate signaling pathways, making it less clear that the primary cause of disease is the associated transcriptomic dysregulation.

### A systematic transcriptomic signature reversal program for neurodevelopmental disorders

The substantial contribution of transcriptomic regulators to genetic neurodevelopmental diseases highlights the enormous potential for developing a transcriptomic reversal program for these diseases. However, this undertaking will require a concerted, community-based effort.

Here, we outline concrete steps and special considerations required to facilitate the successful implementation of this approach.

### Generating mouse and organoid models of disease

Except in rare cases, it is not possible to acquire brain tissue from individuals with neurodevelopmental disorders. Therefore, generating disease expression signatures for these diseases requires robust model systems. Ideal pre-clinical models for this approach need to express the disease-associated gene in the same cell type as *in loco*, recapitulate perturbations to the regulatory network and disease phenotypes, and enable high-throughput screening of compounds^46^. No single model satisfies all of these criteria. For example, *in vitro* models offer the most potential for high-throughput screening and can be developed for human cells, but of course cannot represent developmental or behavioral consequences of neurodevelopmental disease-causing mutations^46^. We therefore suggest an integrated approach that includes both human organoids of brain regions and mouse models.

Genetically engineered mouse lines have contributed to significant advances in neurodevelopmental disease gene research. Importantly, mouse models can provide important behavioral and electrophysiological phenotypic endpoints for small molecule screening^47,48^. Mouse models of genetic epilepsies, for example, may display spontaneous seizures or altered seizure thresholds. Additionally, while mouse models cannot fully capture the entirety of symptoms associated with ASD and schizophrenia, they can model certain aspects of these disorders that may manifest in an equivalent manner in rodents^49^. For example, mouse models of schizophrenia-causing mutations in *SETD1A* and ASD-causing mutations in *CHD8*, display cognitive impairments that overlap with human pathology^50,51^.

Despite the utility of mouse models, there are inherent differences between the development, architecture, and function of the mouse and human brain^52^. Therefore, mouse studies should be compared with models based on human cells that capture as much of human brain development as is currently possible. Cortical organoids, which are self-organizing threedimensional structures that resemble features of human cortical development, have emerged as one such model^53–55^. Although reductionist in nature, this system is amenable to genetic engineering and high-throughput screening. Because organoids are derived from human cells, they also overcome potential species-specific differences in gene expression regulation. In fact, investigators have successfully used organoids to identify molecular phenotypes in various neurodevelopmental conditions that have been difficult to study in mice, such as lissencephaly^56^ and periventricular nodular heterotopia^57^. Organoids can be developed directly using human cells from individuals with the disease of interest, reprogramming those cells into induced pluripotent stem cells (iPSCs), and then differentiating them into cortical organoids. Alternatively, genome-editing can be used to introduce a disease-causing mutation in a control iPSC line. While organoids perhaps bring us closer to human biology, this model system has its own limitations. Most notably, recent work has shown that these artificial systems ectopically activate cellular stress pathways, which can impact cell type specification^58^. Moreover, organoids are now known to show substantial variability from one batch to another, making it challenging to uncover consistent transcriptomic, morphological, or functional phenotypes.

Clearly, neither model is perfect. Mice lack human-specific features of development, while organoids are subject to artificial conditions that impact cellular development. We do not yet know how often gene expression signatures in mouse models will be reflected in organoids of the same model, and vice versa. When mouse and organoid models are congruent, we can be more confident in the relevance of the disease signatures. However, when the expression changes between models differ, we must attempt to elucidate the source of this divergence. For example, if organoids are differentiated with unrelated protocols but retain the same transcriptomic signatures, those signatures are likely relevant to the underlying biology than an experimental artifact. It would be similarly worthwhile to compare transcriptomes of global knockout mice with either targeted knockouts or knock-ins. Both targeted knockouts and knockins should presumably recapitulate the global knockout, while providing granular insight into the functions of the gene of interest. If transcriptomes from global knockouts show important differences compared with targeted mutants however, peripheral tissues may be playing an underappreciated role in the disease biology. Unlike mice, organoids are isolated systems that would miss these effects entirely.

### Deriving disease gene expression signatures via single-cell RNA-sequencing

It is becoming increasingly clear that disease-causing mutations in transcriptomic regulators have cell type-specific effects. One recent study used *in vivo* Perturb-Seq (a pooled CRISPR screening method) to introduce mutations in 35 ASD associated genes within the developing mouse brain *in utero* and then perform scRNA-seq postnatally^59^. Many of the edited genes were transcriptomic regulators, such as *Chd8, Gatad2b, and Set5d*, and perturbations of these genes led to cell type-specific gene expression changes. For example, loss of *Chd8* led to the dysregulation of gene expression programs required for oligodendrocyte differentiation and maturation. In a separate study, scRNA-seq revealed that haploinsufficiency of the schizophrenia-associated gene, *Setd1a*, led to stronger differential gene expression effects in cortical pyramidal cells than in any other cell type^50^. We recently reported that a mouse model of *HNRNPU*-mediated EE results in an increased burden of differentially expressed genes in pyramidal cells of the subiculum^45^. Using CMap, we found that the compounds predicted to reverse the subiculum signature did not strongly overlap with the compounds predicted to reverse the bulk signature^45^.

Together, these studies highlight the need for scRNA-seq in generating cell type-specific disease expression signatures. We argue that each implicated transcriptomic regulator should be modeled and profiled, and the resulting expression data should be made available publicly. Although a massive undertaking, this effort would both maximize the number of transcriptomic reversal successes and equip investigators to identify potentially convergent disease mechanisms among these genes.

### Developing a compendium of compound signatures

Although the CMap provides thousands of compound signatures, most of these signatures were derived from cancer cell lines. A small proportion of compounds (n = 768) were also profiled in iPSC-derived neural progenitor cells (NPCs) and neurons. Using these data, we performed a cluster analysis to compare the differential gene expression responses between neurons, NPCs, and the two most commonly assayed cancer cell lines, MCF7 and PC3. Perhaps unsurprisingly, NPCs and neuron signatures clustered separately from the cancer cell line signatures (**Figure 3A**). Consistent with other reports, these results suggest cell type-dependent gene expression responses^26^. Therefore, compound signatures generated in cancer cell lines may have limited utility in identifying transcriptomic reversal candidates for neurological diseases.

**Figure 3.**
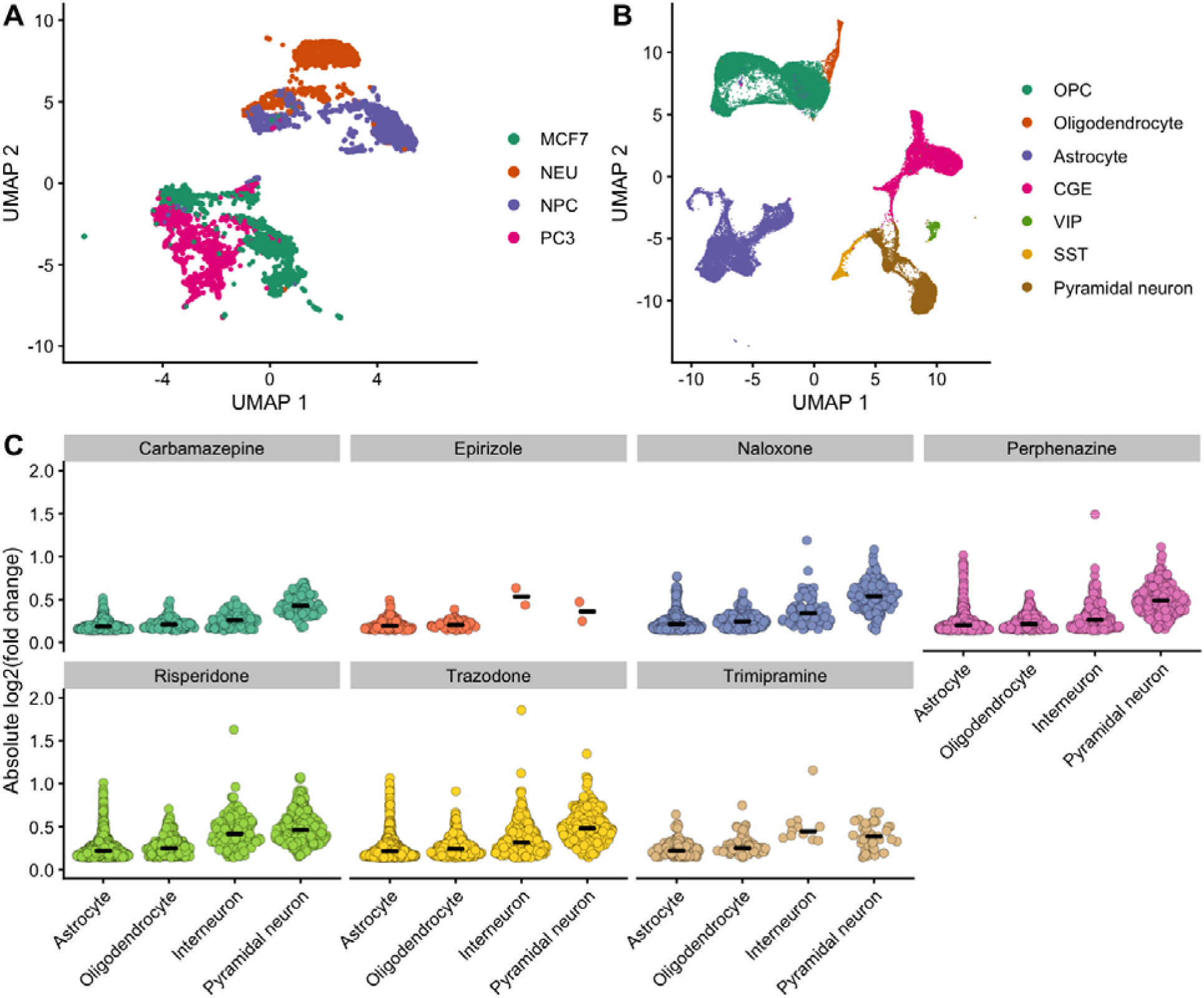
Cell type-specific gene expression responses to small molecules. **(A)** UMAP plot representing differential gene expression responses to roughly 800 small molecules in cancer cell lines (MCF7 and PC3) versus iPSC-derived neurons and neural progenitor cells (NPCs). Gene expression was measured via the L1000 assay. **(B)** UMAP plot representing clusters of major cell types detected via scRNA-seq of cultured primary mouse cortical neurons (47,494 cells). Cells were dosed with one of seven compounds or with DMSO (control). **(C)** Violin plots representing the average effect size per cell type of differentially expressed genes (FDR < .01 and expression change > 10%) in each treatment condition.

Small molecule-induced gene expression changes not only vary between cancer cell lines and neural cells, but also between subtypes of neural cells. To demonstrate this point, we performed single-cell RNA-sequencing of cultured mouse primary cortical neurons treated with seven different FDA-approved compounds with known pharmacological effects on the central nervous system (naloxone, perphenazine, trazodone, trimipramine, risperidone, epirizole, and carbamazepine). These seven compounds were chosen because they elicited the most orthogonal signatures in the CMap NPC data, suggesting they represent the transcriptomic landscape of FDA-approved blood-brain penetrant compounds. We dosed each compound at its reported maximum serum concentration for 24 hours on the 10^th^ day *in vitro* (DIV 10). We then compared expression responses in four major cell types: astrocytes, oligodendrocytes, excitatory neurons, and inhibitory neurons (**Figure 3B**). Interestingly, both the number of differentially expressed genes and magnitude of expression changes varied by markedly cell type and by drug (**Figure 3C**).

The availability of cell type-specific signatures for both drugs and disease models would allow for comparisons in matched cell types. This approach should theoretically reduce the number of false positive transcriptomic reversal candidates. The current per-sample costs of conventional scRNA-seq approaches, however, preclude the generation of even a modestly sized compendium of signatures. Fortunately, emerging multiplexing techniques, such as Cell Hashing^60^, MULTI-seq^61^, and sci-Plex^62^, dramatically reduce costs and increase the feasibility of generating cell type-specific signatures. Ideally, these signatures should be generated in both organoids and cultured primary mouse neurons. Compounds should be dosed at physiologically relevant concentrations, which can be estimated from a compound’s reported maximum serum concentration or maximum therapeutic dose if known.

### Compound prioritization and validation

In any transcriptomic reversal approach, candidate compounds need to be administered to cell**s** containing the mutated gene to verify that they in fact restore the transcriptome toward normal. Once validated, these compounds need to be tested for the functional consequence of thi**s** restoration. We do not yet know to what extent the transcriptome must be normalized to rescu**e** molecular and behavioral phenotypes, and this will only be learned through experience of such studies. Furthermore, it will be crucial to determine whether compounds elicit the same transcriptomic effects *in vivo* as they do *in vitro* through experimental comparisons.

The cell type-specific transcriptomic reversal approach offers unique opportunities during the validation phase. Consider the case in which more than one cell type emerges as particularly vulnerable to a disease-causing mutation. In this instance, it is possible that drugs that reverse the signature in one cell type may not reverse the signature in another. While this scenario would require the interrogation of more compounds, it also offers the ability to probe the contribution of each cell type to the disease phenotype and to identify potential combination therapies. For example, administering compounds that revert the transcriptome in one cell type may rescue some phenotypes, whereas compounds that revert the transcriptome of another cell type could rescue others. Finally, the prioritization of compounds should also consider the safety profile of disease-signature reversing compounds. Ideally, a compound would reverse the disease signature while having minimal impact on the rest of the transcriptome. In reality, this ideal is unlikely to be often achievable. Approaches will need to be developed to assess whether expression changes beyond the disease-signature reversal are likely to be harmful. For example, one potential approach would be to consider how many genes that are genetically intolerant^63–65^ and that are intolerant to expression variation^66^ have been perturbed.

## Conclusions

We have outlined a precision medicine paradigm that could lead to the discovery of numerous treatment options for neurodevelopmental disorders caused by mutations in transcriptomic regulators. We argue that the paradigm can be most easily refined by working on these Mendelian conditions, in which we can be confident of the causal consequences of transcriptomic alterations. Once the key aspects of this paradigm are better understood in the context of neurological disorders, such as how much restoration of the altered transcriptome is required, it will be possible to extend the paradigm to conditions with more complex genetics, such as the more common forms of the aforementioned diseases.

Establishing such a systematic program would require a community-based effort and substantial data sharing. As demonstrated by the CMap, publicly accessible drug signatures can open the floodgates for identifying novel therapies. Furthermore, meeting the goals outlined here, including the establishment of robust disease models, implementation of large-scale scRNA-seq analyses, and the evaluation of candidate compounds, will require the collaborative efforts between researchers with biological, clinical, and computational expertise. Given the remarkable importance of genes with transcriptomic effects in neurodevelopmental disorders we believe that careful implementation of a paradigm of transcriptomic-reversal has the potential to identify effective treatments for many genetic diseases.

## Methods

### Connectivity Map analysis

We compared Connectivity Map^17^ L1000 data for 2,103 compounds that were dosed in MCF7 cells, PC3 cells, iPSC-derived neurons, and iPSC-derived neural progenitor cells (NPCs). Data were downloaded from the Gene Expression Omnibus (accession number GSE92742). We only considered gene expression values for the 978 genes directly assessed by the L1000 assay. We then created a UMAP plot to visualize clustering patterns for gene expression signatures elicited by these 2,103 compounds.

To find compounds that elicited orthogonal gene expression signatures in neural cells, we considered the L1000 data for blood-brain barrier penetrant compounds that were assayed in NPCs. We opted to use the NPC signatures rather than neuron signatures as there were more compounds assayed in NPCs (141 versus 53). We performed clustering using affinity propagation^67^, which resulted in 7 clusters with the following examplars (i.e. cluster members that are representative of the group): naloxone, perphenazine, trazodone, trimipramine, risperidone, epirizole, and carbamazepine. These compounds were chosen for dosing in mouse cortical neurons.

### Mouse husbandry

Experiments were performed on the inbred background C57BL/6NJ (005304 JAX stock). All mice were maintained in ventilated cages with controlled humidity at ~60%, 12h:12h light:dark cycles (lights on at 7:00AM, off 7:00PM) and controlled temperature of 22–23°C. Mice had access to regular chow and water, ad libitum. Breeding cages were fed a high fat breeder chow. Mice were maintained and all procedures were performed within the Columbia University Institute of Comparative Medicine, which is fully accredited by the Association for Assessment and Accreditation of Laboratory Animal Care. All protocols were approved by the Columbia Institutional Animal Care and Use Committee.

### Primary cortical culture

Prior to use, 24-well TC Treated plates (Corning) were coated overnight with 50 μg/mL poly-D-lysine (Sigma) in 0.1M borate buffer (pH 8.5). Cortices acquired from multiple sets of male littermates were incubated in activated 20 U/mL Papain/DNase (Worthington) for 15 minutes at 37°C, centrifuged at 300 x g for 5 min, and washed in 1X PBS. Cell pellets were suspended in NBA/B27 [consisting of Neurobasal-A (Life Technologies), 1X B27 (Life Technologies), 1X GlutaMax (Life Technologies), 1% HEPES, and 1X Penicillin/Streptomycin], supplemented with 1% fetal bovine serum (Gibco) and 5 μg/mL laminin. Five hundred thousand cells were plated per well. The day after plating, media was removed and replaced with pre-warmed NBA/B27. Cultures were maintained at 37°C in 5% CO_2_. 50% of the medium was changed every other day with NBA/B27 starting on DIV3. On DIV10, seven compounds and the control media were introduced to the appropriate wells in duplicate at their reported maximum serum concentrations^68^:

| Drug | Maximum serum concentration (nM) | Dose (nM) |
| --- | --- | --- |
| Naloxone | 91.7 | 100 |
| Perphenazine | 49.4 | 50 |
| Trazodone hydrochloride | 4302.7 | 4000 |
| Trimipramine maleate salt | 1018.9 | 1000 |
| Risperidone | 48.7 | 50 |
| Epirizole | Unknown | 1000 |
| Carbamazepine | 84.6 | 100 |
| DMSO (negative control) | - | - |

After 24 hours of compound exposure, the cells were dissociated with activated 20 U/mL Papain, strained through 40 um cell strainers (MTC Bio) to remove cell clumps and collected for single-cell RNA-sequencing (scRNA-seq) on the 10X Chromium in two batches. scRNA-seq libraries were constructed using the 10X Chromium Single Cell 3’ Reagent Kits v2 according to manufacturer descriptions, and samples were sequenced on a NovaSeq 6000. Reads were aligned to the mm10 genome using the 10X CellRanger pipeline with default parameters to generate the feature-barcode matrix.

### scRNA-seq analysis

We used Seurat v3 to perform downstream QC and analyses on feature-barcode matrices^37,38^. We removed all genes that were not detected in at least 4 cells. We further removed cells with fewer than 200 genes or more than 2,500 genes detected. We further removed all cells with greater than 25% of reads mapping to mitochondrial genes. 47,494 cells remained after filtering:

| Drug | Batch 1 | Batch 2 | Total |
| --- | --- | --- | --- |
| Naloxone | 2203 | 3360 | 5563 |
| Perphenazine | 2457 | 4254 | 6711 |
| Trazodone | 2298 | 3482 | 5780 |
| Trimipramine | 3516 | 3299 | 6815 |
| Risperidone | 2367 | 2757 | 5124 |
| Epirizole | 3051 | 2982 | 6033 |
| Carbamazepine | 3189 | 3021 | 6210 |
| DMSO (control) | 1770 | 3488 | 5258 |

The filtered matrices were log-normalized and scaled to 10,000 transcripts per cell. We used the variance-stabilizing transformation implemented in the FindVariableFeatures function in order to identify the top 2,000 most variable genes per sample. We used Seurat’s data integration method to harmonize gene expression across datasets prior to clustering. We first identified anchors between samples in each dataset using the FindIntegrationAnchors function, which uses canonical correlation analysis (CCA) to identify pairwise cell correspondences between samples. We then computed an integrated expression matrix using these anchors as input to the IntegrateData function.

Next, we used linear regression to regress out the number of UMIs per cell and percentage of mitochondrial reads using the ScaleData function on the integrated expression matrices. We then performed dimensionality reduction using PCA. For each dataset, we selected the top 30 dimensions to compute a cellular distance matrix, which was used to generate a K-nearest neighbor graph. The KNN was used as input to the Louvain Clustering algorithm implemented in the FindClusters function. For clustering via Louvain, we chose a resolution parameter of 0.8. We visualized the cells using UMAP via the RunUMAP function. To annotate and merge clusters, we performed differential gene expression analysis on the integrated expression values between each cluster using the default parameters in the FindMarkers function, which implements a Wilcoxon test and corrects p-values via Bonferroni correction. Additionally, we visualized the expression of canonical marker genes aggregated from previous single-cell publications^71-74^.

### Differential gene expression analysis

We performed cell-type-specific differential gene expression analysis using MAST^75^, as implemented in Seurat’s FindMarkers function, in order to identify genes dysregulated between drug-treated and DMSO-treated cells. We excluded all non-coding genes, genes encoding ribosomal proteins, and pseudogenes from our analysis to reduce the multiple testing burden. For each cell type, we fit a linear mixed model that included the gene detection rate (ngeneson) and sequencing batch as latent variables. We corrected the p-values using the Benjamini-Hochberg FDR method, and considered genes with an FDR < 0.01 as differentially expressed.

## Acknowledgments

We are grateful to Wayne Frankel for helpful discussions and his comments on the manuscript.

